# Duchenne Muscular Dystrophy Cell Culture Models Created By CRISPR/Cas 9 Gene Editing And Their Application To Drug Screening

**DOI:** 10.1101/2020.02.24.962316

**Authors:** P. Soblechero-Martín, E. Albiasu-Arteta, A. Anton-Martinez, L. de la Puente-Ovejero, I. Garcia-Jimenez, G. González-Iglesias, I. Larrañaga-Aiestaran, A. López-Martinez, J. Poyatos-García, E. Ruiz-Del-Yerro, F. Gonzalez, V. Arechavala-Gomeza

**Author notes:** **Address for correspondence:** Prof. Virginia Arechavala-Gomeza, Biocruces Bizkaia Health Research Institute, Barakaldo, Bizkaia, 48903, Spain.

## Abstract

Gene edition methods are an attractive putative therapeutic option for Duchenne muscular dystrophy and they have an immediate application in the generation of research models. To generate two new edited myoblast cultures that could be useful *in vitro* drug screening, we have optimised a CRISPR/Cas9 gene edition protocol. We have successfully used it in wild type immortalised myoblasts to delete exon 52 of the dystrophin gene: DMDΔ52-Model, modelling a common Duchenne muscular dystrophy mutation; and in patient’s immortalised cultures we have deleted an inhibitory microRNA target region of the utrophin UTR, leading to utrophin upregulation. We have characterised these cultures and, to show their use in the assessment of DMD treatments, we have performed exon skipping in the DMDΔ52-Model and have used the unedited cultures/ DMD-UTRN-Model combo to assess utrophin overexpression after drug treatment. While the practical use of DMDΔ52-Model is limited to the validation to our gene edition protocol, DMD-UTRN-Model offers a possible therapeutic gene edition target as well as a useful positive control in the screening of utrophin overexpression drugs.

## INTRODUCTION

### Duchenne muscular dystrophy

(DMD) is a fatal X-linked recessive disease affecting one out of 5.000 newborn males. It is commonly caused by deletions disrupting the open reading frame of the DMD gene causing a lack of dystrophin protein^1^. Patients carrying out of frame mutations present a severe phenotype, while those carrying in-frame mutations may result in hypomorphic alleles, a partially functional dystrophin and milder phenotypes, such as in Becker muscular dystrophy (BMD)^2^. Dystrophin plays a major role in membrane stabilization during muscle contraction, linking the actin cytoskeleton to the sarcolemma^3^ and also contributes to extracellular signalling^4^. Lack of dystrophin in DMD patients’ muscles leads to progressive muscle wasting and degeneration. DMD children suffer from loss of ambulation in the first or second decade of life and premature death by cardiac and respiratory complications^5^.

Although no definitive cure for DMD is available, four drugs have been recently approved by different regulatory agencies: ataluren, approved by the European Medicines Agency (EMA), induces readthrough of premature stop codons during mRNA translation, generating a full length dystrophin protein^6^; and the US Food and Drug Administration (FDA), has recently approved three antisense oligonucleotide drugs, which modulate splicing by exon skipping, restoring DMD reading frame and leading to a shorter but functional protein^7,8^: eteplirsen, golodirsen and viltolarsen. Exon skipping therapies aim to attenuate the phenotype and phenocopy milder BMD-like genotypes, to potentially improve disease outcomes. Eteplirsen targets DMD exon 51 and golodirsen and viltolarsen, exon 53. All approved drugs are mutation specific and designed to rescue specific patient mutations only present, respectively, in 13% (ataluren), 13% (eteplirsen) and 8% (golodirsen and viltolarsen) of patients^9^. It is therefore important to assess exon-skipping strategies targeting other DMD exons, and many such drugs are in different phases of clinical assays^10^.

Other than these approved therapies, gene transfer has emerged as a promising approach applicable to all DMD patients. SRP-9001, a human micro-dystrophin gene transfer therapy, driven by the muscle-specific promoter MHCK7 and delivered by the recombinant adeno-associated virus rAAVrh74 has recently got the Fast Track designation by the FDA after showing a robust posttreatment increase of micro-dystrophin expression in patients muscles and positive safety and tolerability data in clinical trials^11^ suggesting the potential of this therapy to provide clinically meaningful functional improvement in DMD patients.

Alternatively to these therapies aiming to restore dystrophin expression, many compounds targeting secondary effects of dystrophin deficiency or looking for alternatives to dystrophin are also under evaluation.

*In vitro* cellular models are particularly useful to assess the efficiency of novel therapies for DMD. However, only a few human immortalized muscle cell lines derived from DMD patients are currently available^12^. Due to the wide spectrum of DMD mutations and the difficulties to obtain DMD patient muscle biopsies, DMD-myoblasts models would provide a powerful resource for *in vitro* drug screening and study disease rescue mechanisms.

### UTROPHIN

*(UTRN)* is an autosomal paralog of dystrophin, expressed in skeletal muscle cells during embryonic development, but restricted to neuromuscular and myotendinous junctions in the mature muscle fibre^13^. Overexpression of utrophin in skeletal muscle in DMD animal models can partially compensate the lack of dystrophin and improve DMD phenotype^14–16^. Importantly, ectopic and high levels of utrophin in myoblasts are not associated with toxicity, making utrophin upregulation an interesting therapeutic strategy applicable to all patients, regardless of their specific mutation^17,18^. Ezutromid/SMT-C1100 was the first utrophin modulator evaluated in clinical assays, but was recently abandoned due to lack of evidence of utrophin restoration, nor clinical improvement demonstrated in patients ^19,20^. Alternatively, other studies proposed new strategies to upregulate *UTRN* by blocking the inhibitory target region of microRNAs repressing *UTRN* expression^21^. Recently, utrophin upregulation has been efficiently achieved using gene therapy in multiple animal models with non-immunogenic side effects^22^. Several new compounds that aim to overexpress utrophin are currently being developed^23,24^, and this preclinical development could benefit from a gold standard or an adequate positive control to use in these assays.

### CRISPR/Cas9

currently represents a very efficient and versatile genome-engineering tool, introducing small and large DNA modifications, including large genomic deletions in different cell types and organisms^25^. In the presence of two single guide (sg) RNAs targeting two different loci on the same chromosome, Cas9 can induce two DNA double strand breaks (DDSBs), leading in some cases to deletion of the excised DNA segment through repair by the non-homologous end joining (NHEJ) pathway^26^. Similar to antisense oligo-mediated exon skipping therapies at RNA level, CRISPR/Cas9 can therefore be used to remove mutations by deleting mutated exons and restore the open reading frame of the *DMD* gene^27–29^. The advantage of this approach is that the genetic modification, once introduced, is stable over cell cycles. CRISPR/Cas9 has been successfully employed to correct mutations and/or restore the open reading frame recovering dystrophin expression both *in vitro* and *in vivo*^30,31^. However, some hurdles have been reported, such as difficulties transfecting myoblast or a recent study in the golden retriever muscular dystrophy dog (GRMD), where no dystrophin restoration at protein level was evident after gene editing using this technology^32^. More importantly, as well as these preclinical problems others such as possible off-target problems or immunogenicity linked with the use of Cas9^33^ may delay their clinical application. However, while gene editing as therapeutic option still needs further development, CRISPR/Cas9 methodology has been applied to provide a large number of new animal models to further understand DMD pathology and perform preclinical studies^34^. In our quest to optimise the preclinical development of new therapies for DMD, we have developed a genome editing strategy applicable in control and patient myoblasts. Here, we report two new cell cultured models that can be successfully used for preclinical assessment of new DMD therapies: a culture that replicates a patient’s deletion (**DMDΔ52-Model**) and another that overexpresses utrophin (**DMD-UTRN-Model**).

## RESULTS

### Generation of cell culture models by gene editing

We completed two different gene editing projects: **objective 1** aimed to delete exon 52 of the DMD gene (a common mutation in *DMD* patients) in control immortalised human myoblasts to generate a disease model (**DMDΔ52-Model**); **objective 2** (**DMD-UTRN-Model**) was to delete in the UTRN gene of DMD immortalized human myoblasts an inhibitory microRNA target region to generate a utrophin ectopic expression rescue model.

Each edition required to design two sgRNAs flanking the region to be deleted to generate two DDSBs leading to the removal of that region (Figure 1A and D). All the 25 different combinations of the sgRNAs designed, cutting before (x5) and after (x5) the target region, were tested in HEK293 cultures first (Supplementary figure 1A and B). The most efficient combination of sgRNAs in HEK293 cells for each objective was selected to be used in the transfection of human immortalized myoblasts (Supplementary figure 1C and D).

**Figure 1.**
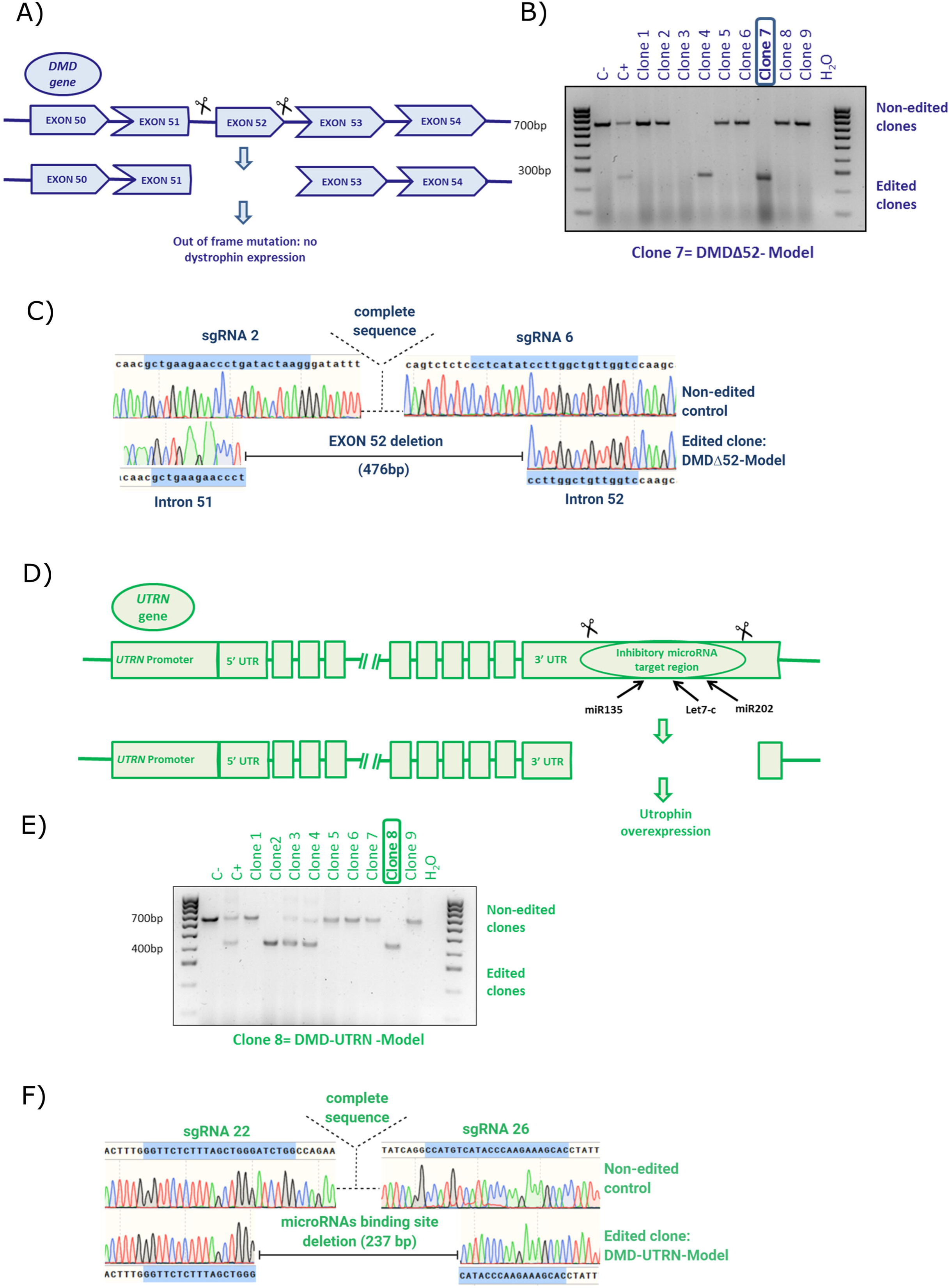
Editing approach and genotyping *DMD* and *UTRN* deletion breakpoints in edited myoblast clones. (A and D). Schematic representation of our strategy for editing the *DMD* (A) and the *UTRN* loci (D). A pair of flanking sgRNAs were co-transfected in order to delete *DMD* exon 52 (A) or the inhibitory microRNA target region contained in the 5’UTR of *UTRN* (D). (B and C) PCR genotyping of *DMD* edited clones (B) and Sanger sequencing of clone 7 or DMDΔ52-Model (C). (E and F) PCR genotyping of *UTRN* edited clones (E) and Sanger sequencing of clone 8 or DMD-UTRN-Model (F). Larger products in agarose gels (B and E) indicate non-edited clones, and shorter ones correspond with the expected deletion. (C and F) Sequences of the smaller bands confirmed the expected gene editing for objective 1: DMDΔ52-Model (C) and objective 2: DMD-UTRN-Model (F).

To accomplish objective 1, two GFP-plasmids, each encoding Cas9 nuclease and either sgRNA 2 or sgRNA 6, were transfected into human immortalized control myoblasts. For objective 2, the selected GFP-plasmids, encoded Cas9 nuclease and either sgRNA 22 or sgRNA 26 and were transfected into human immortalized DMD myoblasts. After FACS sorting of individual GFP-positive cells, clones were expanded for DNA extraction (Supplementary figure 2). Clones were analysed to confirm the presence of the desired deletions by genomic PCR performed with specific primers for each targeted gene (Figure 1B and 1E), amplicons corresponding in size with the expected deletions were analysed by Sanger sequencing, and the expected deletions were confirmed in all the positive clones (Figure 1C and 1F).

For **objective 1** two positive clones were obtained, clone number 2 and 7 but only clone 7 was used for further analysis and called **DMDΔ52-Model** (Figure 1B and 1C). For **objective 2**, two clones were edited in only one allele, corresponding to numbers 3 and 4 and other two were completely edited, numbers 2 and 8. In this case, clone number 8 was selected to be used for further analysis and was called **DMD-UTRN-Model** (Figure 1E and 1F).

To evaluate any potential off-target effects, each selected sgRNA was analysed *in silico* using the bioinformatics web-tool CRISPOR^35^. We selected the six more likely off-target sites for each sgRNA and analysed each one of them through PCR, followed by Sanger sequencing in edited clones (Table 2 and 3). We found no off-target effects in any of the 12 sites studied for each clone sites (Supplementary figure 3).

**Table 1.**
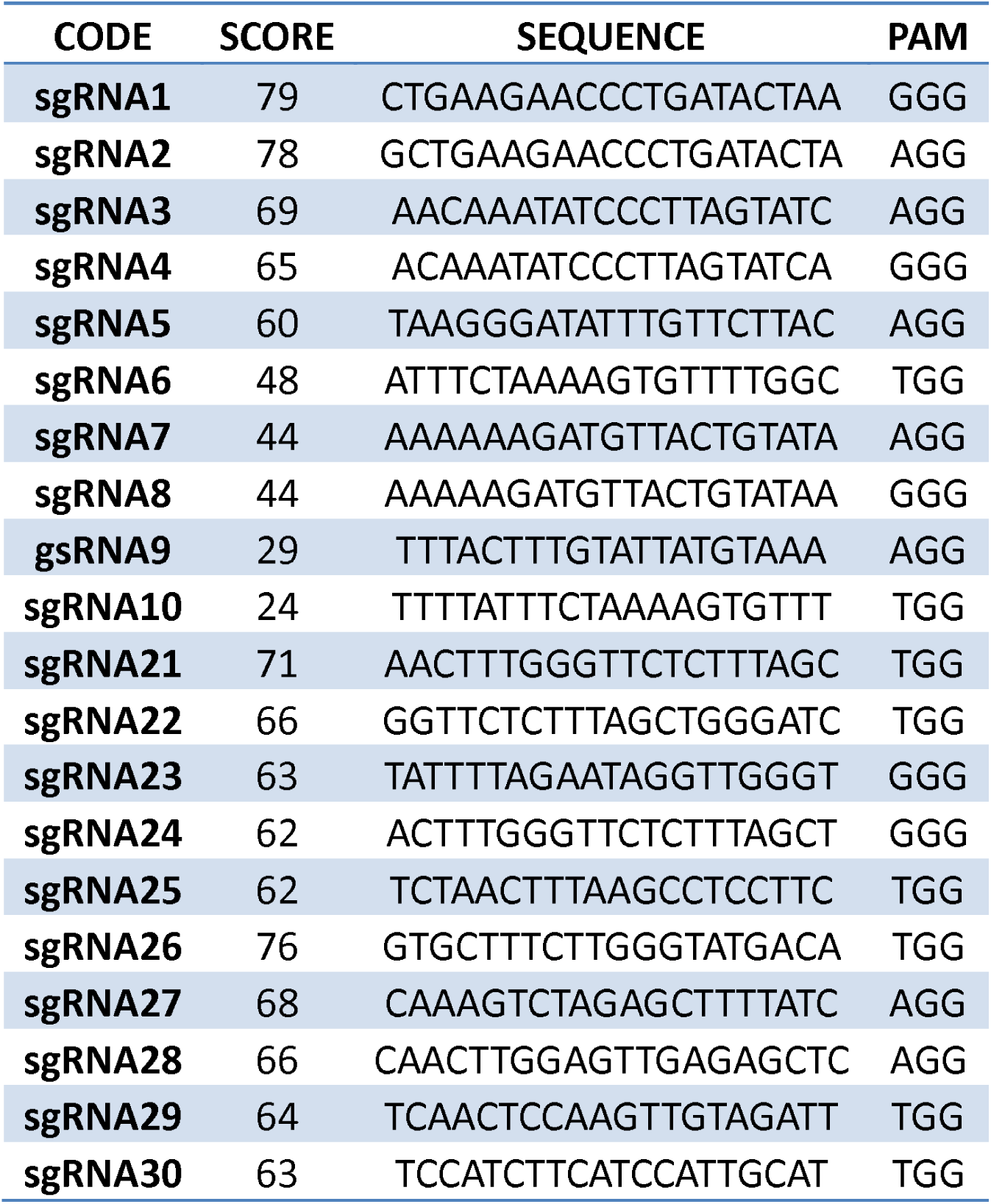
List of sgRNAs. List of sgRNAs for editing the *DMD* and the *UTRN* loci, showing sgRNAs sequences, PAM sequences and scores of all sgRNA tested.

**Table 2.**
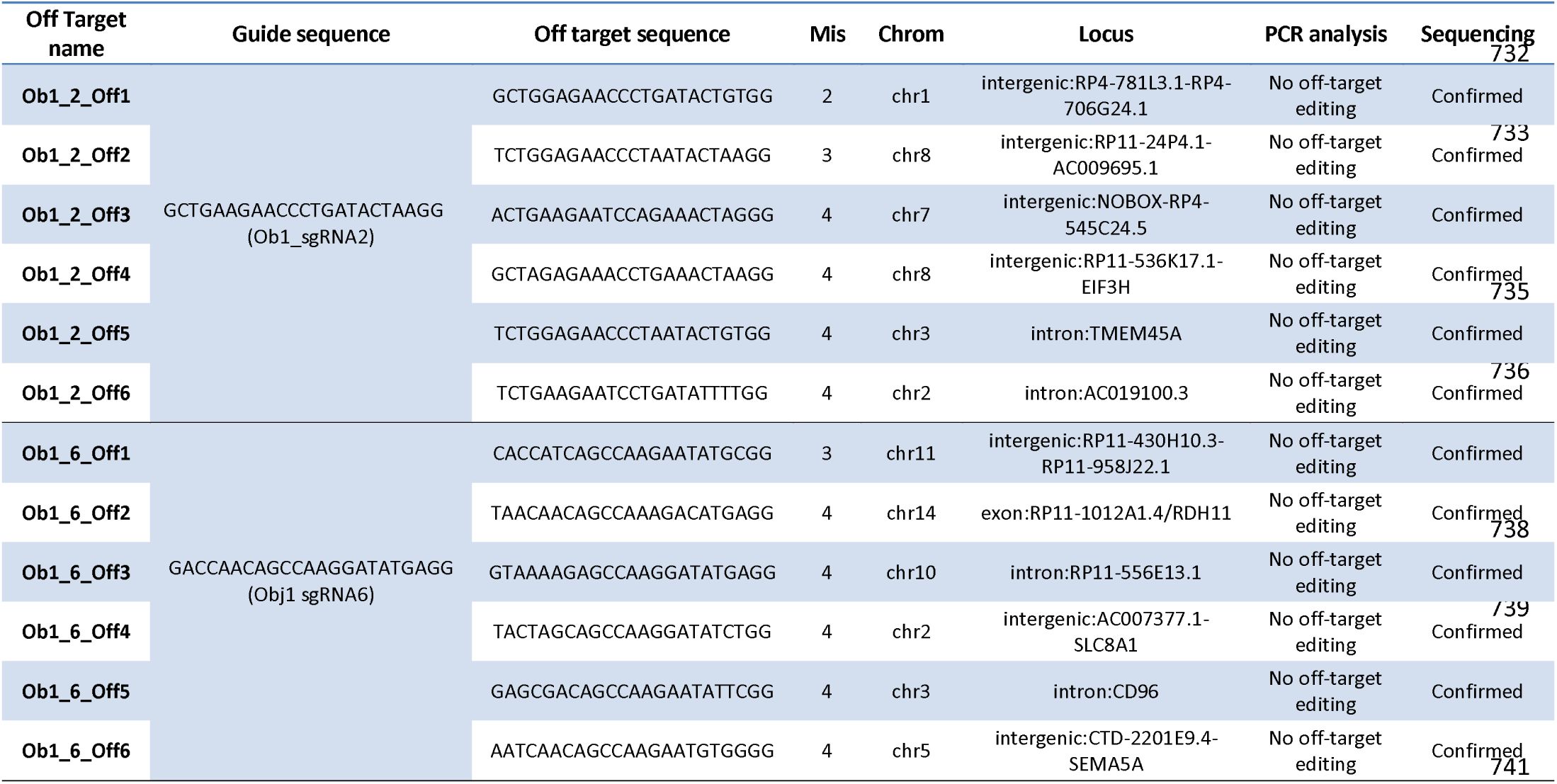
Off-target sites analyses. Top 6 off-target sequences of Ob1sgRNA2 and Ob1sgRNA6 identified with CRISPOR webtool, including the mismatches between sgRNAs, the off-target sequence, the chromosomes and loci targeted. All of them were analysed by PCR and Sanger sequencing, and no off-targets were detected.

**Table 3.**
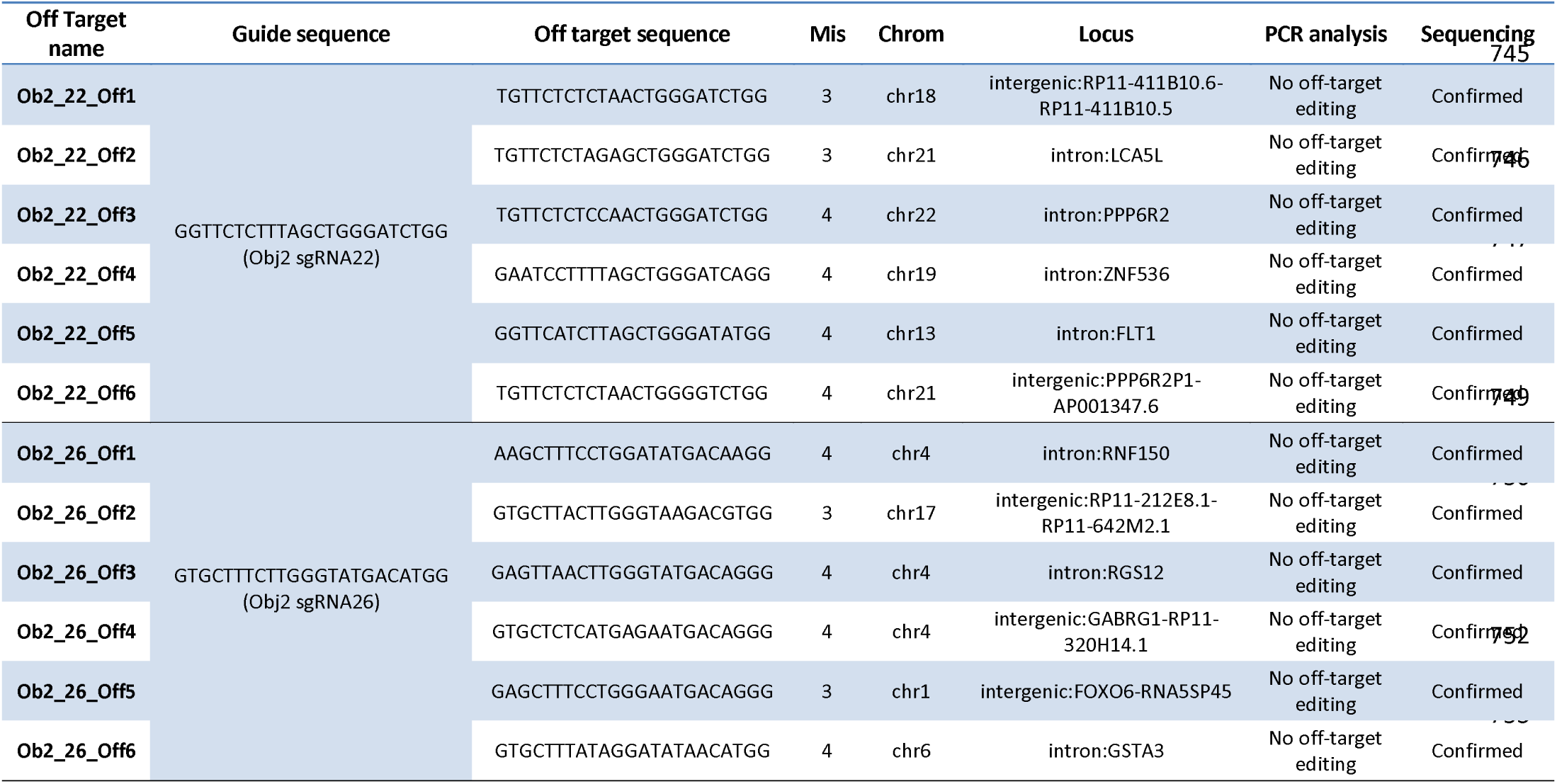
Off-target sites analyses. Top 6 off-target sequences of Ob2sgRNA22 and Ob2_sgRNA26 identified with CRISPOR webtool, including the mismatches between sgRNAs, the off-target sequence, the chromosomes and loci targeted. All of them were analysed by PCR and Sanger sequencing, and no off-targets were detected.

### Analysis of dystrophin and utrophin expression in edited clones

We compared dystrophin expression in myotubes of the DMDΔ52-Model to controls and DMD cultures and confirmed that it was abolished by immunocytochemistry (Figure 2A), western blot analysis (Figure 2B), myoblots (Figure 2C) and droplet digital PCR (ddPCR) (Figure 2D). Dystrophin levels in this model, where exon 52 had been removed by CRISPR/Cas9 editing, were statistically no different than those seen in a culture from a DMD patient.

**Figure 2.**
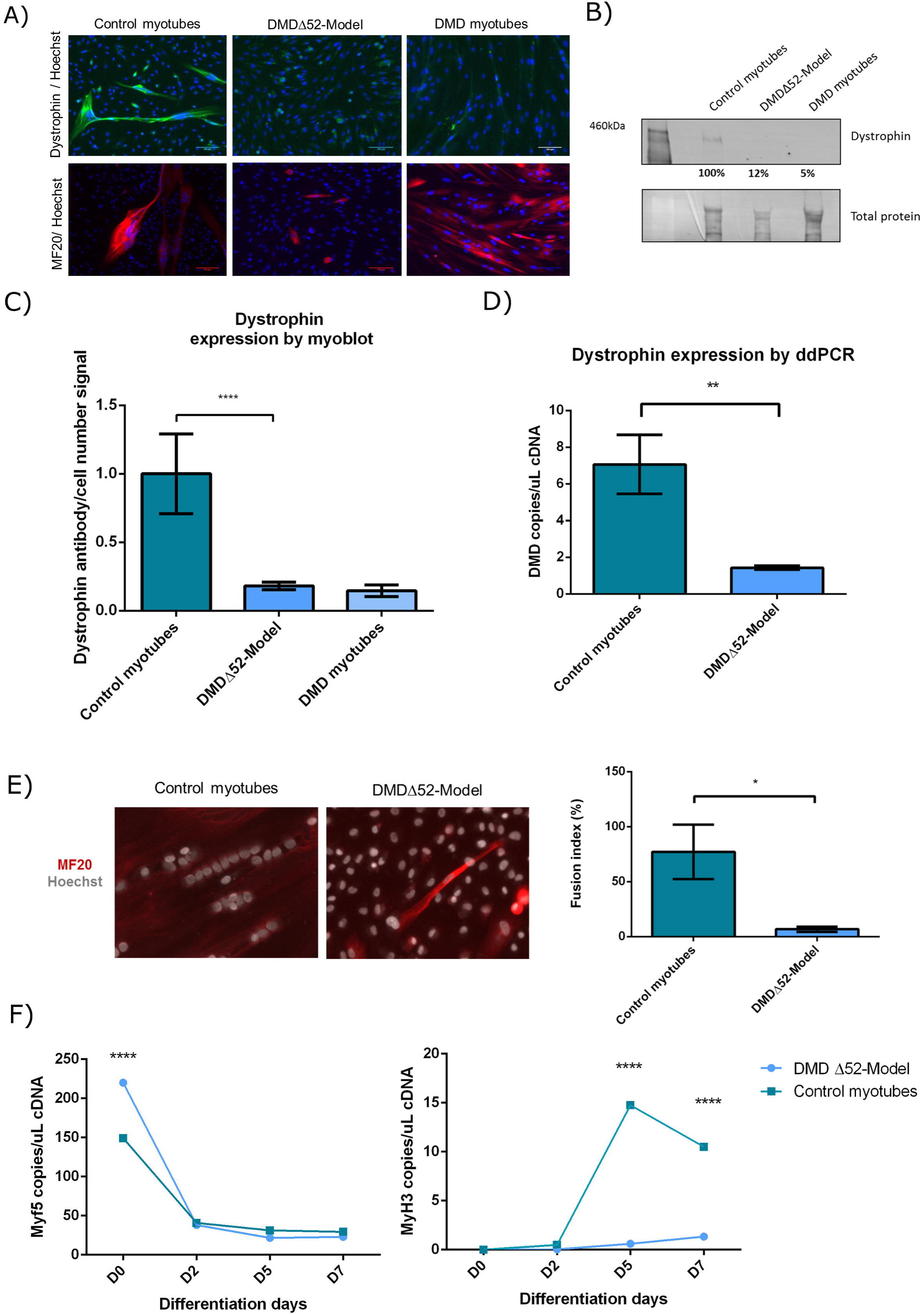
Characterization of DMDΔ52-Model cultures. Dystrophin expression in DMDΔ52-Model cultures compared to control myotubes and DMD myotubes were studied by immunocytochemistry (A), western blotting (B) and myoblots (C). Myoblot experiments, where n=24 wells per cell type were compared, were performed twice. (D) Dystrophin expression in DMDΔ52-Model cultures was compared to control myotubes by ddPCR. For ddPCR experiments three technical replicates per sample and condition were run in parallel and a no template control (NTC) was included as negative control. (E) Differentiated myotubes of DMDΔ52-Model and control cultures were inmunostained with MF20 and Hoechst antibodies. Fusion index was calculated as the ratio between the number of nuclei in differentiated myotubes (defined as >2 nuclei and MF20-positive cells) compared to the total number of nuclei. For quantification, five fields per cell line were randomly chosen and more than 200 nuclei were counted. Analysis was performed using ImageJ software. (F) Differentiation markers, Myf5 and MyH3, were studied by ddPCR at different fusion times in DMDΔ52-Model cultures compared to control myotubes. For ddPCR experiments three technical replicates per sample and condition were run in parallel and a no template control (NTC) was included as negative control. (*p-value <0.05, ** p-value <0.01, ****p-value<0.0001). (P values were determined with Mann-Whitney U test).

Immunocytochemistry showed the increase of utrophin expression between unedited DMD and DMD-UTRN-Model myotubes (Figure 3A) and this increase was corroborated by western blot (a 175% increase, Figure 3B), myoblot analysis (close to 50% increase, Figure 3C) and ddPCR (Figure 3D).

**Figure 3.**
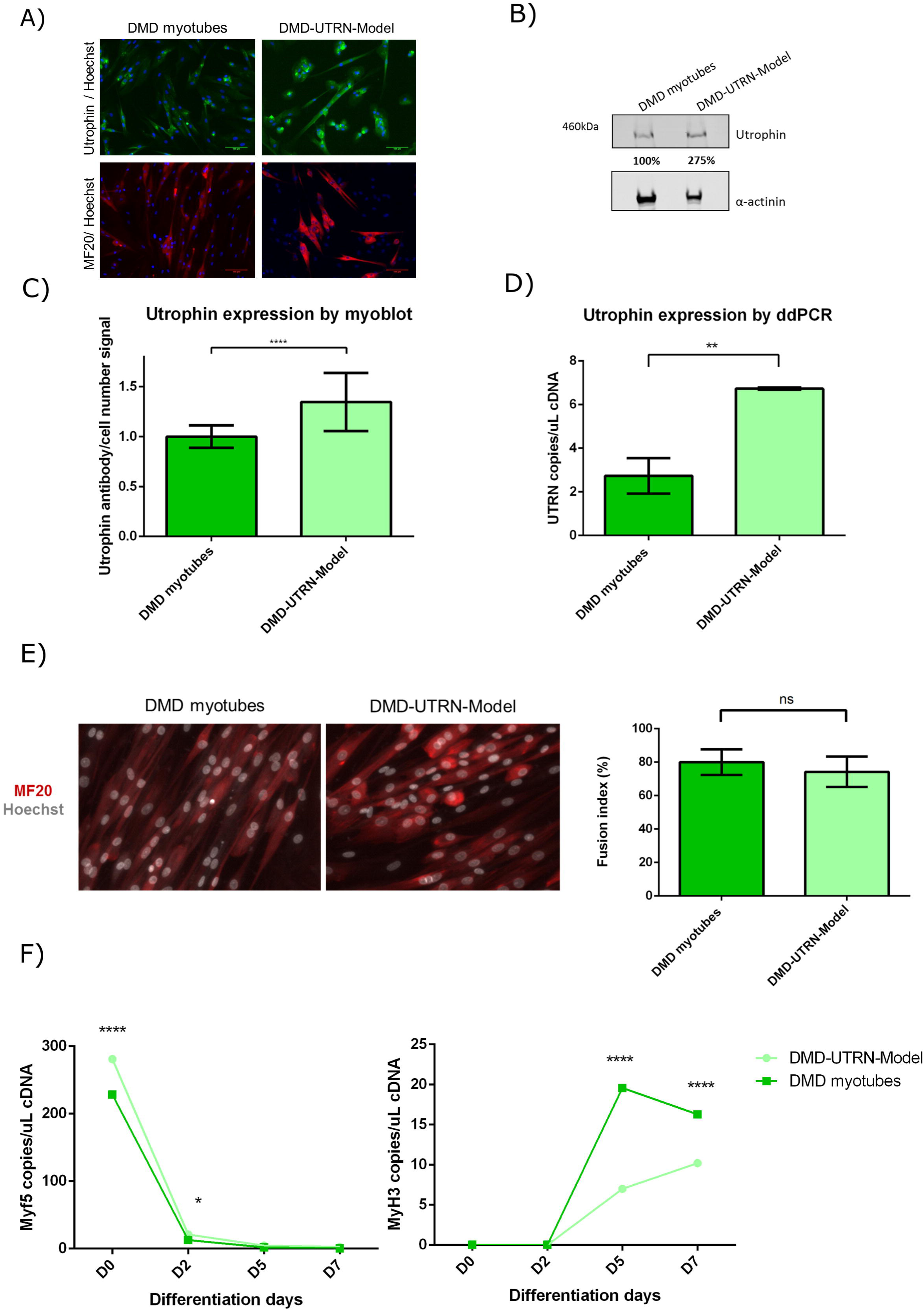
Characterization of DMD-UTRN-Model cultures. Utrophin expression in DMD myotubes compared to DMD-UTRN-Model studied by immunocytochemistry (A), western blotting (B) and myoblots (C). Myoblot experiments, where n=48 wells per cell type were compared, were performed twice. (D) Utrophin expression in DMD-UTRN-Model cultures was compared to DMD myotubes by ddPCR. For ddPCR experiments three technical replicates per sample and condition were run in parallel and a no template control (NTC) was included as negative control. (E) Differentiated myotubes of DMD-UTRN-Model and DMD cultures were inmunostained for MF20 and Hoechst. Fusion index was calculated as the ratio between the number of nuclei in differentiated myotubes (defined as >2 nuclei and MF20-positive cells) compared to the total number of nuclei. For quantification, five fields per cell line were randomly chosen and more than 200 nuclei were counted. Analysis was performed using ImageJ software. (F) Differentiation markers, Myf5 and MyH3, were studied by ddPCR at different fusion times in DMD-UTRN-Model cultures compared to DMD myotubes. For ddPCR experiments three technical replicates per sample and condition were run in parallel and a no template control (NTC) was included as negative control. (*p-value <0.05, ** p-value <0.01, ****p-value<0.0001). (P values were determined with Mann-Whitney U test).

We also quantified dystrophin and utrophin expression in the edited cultures by myoblot during their differentiation process and compared these with control and DMD myotubes (Supplementary figure 4, bar chart).

### Analysis of differentiation markers expression in edited clones

As we suspected that the editing and cloning process could have affected the differentiation of the edited models, the fusion index (%) of edited and non-edited myotubes after MF20 and Hoechst inmunocytochemistry was calculated (Figure 2E and 3E) and the expression of different myogenic regulatory factors like the myogenic factor 5 (Myf5) and the myosin heavy chain isoform 3 (MyH3) were analysed at different time points during myotube formation in DMDΔ52-Model and DMD-UTRN-Model as well as in their corresponding unedited cultures by ddPCR (Figure 2F and 3F). Fusion index was clearly lower in DMDΔ52-Model compared to control myoblasts (Figure 2E) while no significant differences were found between DMD-UTRN-Model and DMD cultures (Figure 3E). As expected, Myf5 expression decreased while MyH3 increased during the differentiation process in both models and followed the same pattern in their origin cells, however we see that MyH3 marker is significantly lower in the edited models at day 5 and day 7 after starting fusion (Figure 2F and 3F).

The MF20 differentiation marker was also analysed by myoblot in edited cultures and we could observe a decrease in both edited clones, no matter the deletion, compared with their corresponding controls (Supplementary figure 4, red lines).

### Evaluation of therapies in newly generated model cell lines

To assess if the DMDΔ52-Model cell culture could be useful to test potential mutation specific therapies for DMD, we evaluated the exon skipping efficiency of an antisense oligonucleotide in this culture. We treated the DMDΔ52-Model cultures with an antisense oligonucleotide drug that can skip exon 51^36^ and restore *DMD* open reading frame. After treatment with this drug, we confirmed that exon skipping had taken place at RNA level (Figure 4A), and a limited restoration of dystrophin expression by myoblot analysis (Figure 4B).

**Figure 4.**
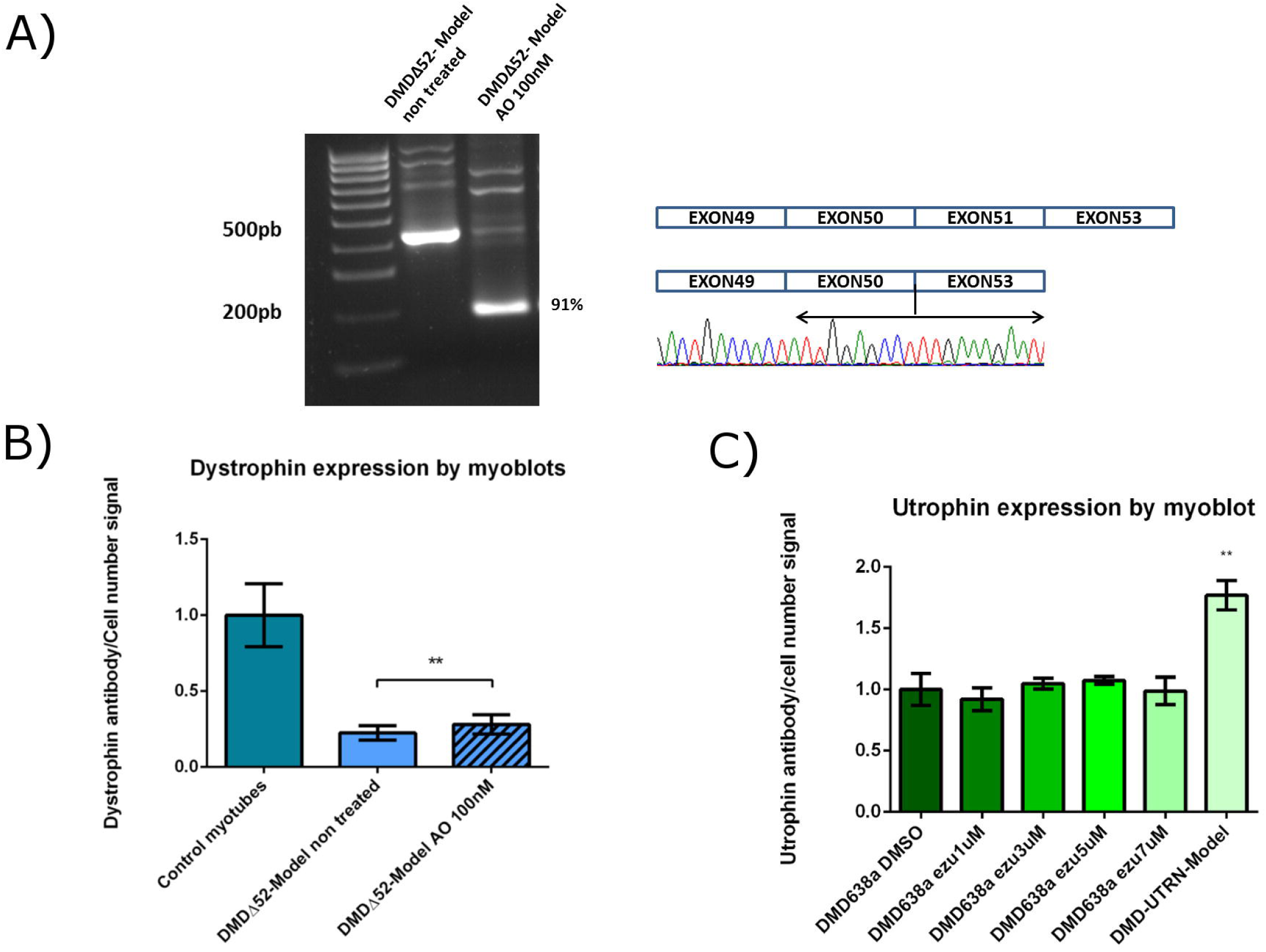
Evaluation of potential therapies in the generated cell culture models. (A) DMDΔ52-Model cell line was treated with an antisense to skip exon 51 and the effect was evaluated by RT-PCR and nested PCR analysis. Gel picture shows a pattern corresponding with the correct skipping, which was confirmed by Sanger sequencing. The same experiment evaluated by myoblot using n=20 replicate wells per condition (B) showed the restoration of dystrophin expression in the treated cultures compared to non-treated (*P values<0.05. P values were determined with Mann-Whitney U test). (C) The DMD-UTRN-Model was used as a positive control in an experiment in which unedited DMD cultures were treated with different ezutromid concentrations to up-regulate utrophin expression. Myoblot analysis using n=8 replicate wells per condition of the treated cultures shows that ezutromid had no significant effect in DMD cultures while utrophin expression is significantly increased in DMD-UTRN-Model compared to unedited DMD cultures. (**P values <0.01. P values were determined with Mann-Whitney U test).

To test our DMD-UTRN-Model as a positive utrophin overexpression control, we cultured it alongside the original unedited DMD cultures, which we treated with several concentrations of ezutromid and we evaluated the expression of utrophin in all cultures. We observed that utrophin was hardly modified in DMD cultures treated with ezutromid while a robust overexpression was confirmed in the DMD-UTRN-Model compared to the unedited DMD cultures (Figure 4C). We also confirmed that this overexpression was stable during a time course experiment (Supplementary figure 4D).

## DISCUSSION

CRISPR/Cas9 has been explored in the past as a possible therapeutic approach for DMD in *in vitro* models of the disease. In most of these cases, a “permanent” exon skipping approach was selected where a shorter protein would be produced^27,28^, while in some others full length dystrophin was the result of the edition^37–39^. Several studies have shown efficacy in mice models^40–42^, an most recently in dogs, which is currently the most advanced example of its application to DMD^29^ but there are also examples of the challenges associated to gene editing^32^. Before this can be a credible therapeutic option, several delivery and manufacturing problems will need to be overcome. In the meantime, this methodology is very useful for researchers looking for cell culture models: muscle biopsies are not routinely collected during diagnosis of this disorder and seldom cultured. This means there are few good culture models of the disease. Another tool created to facilitate research was the immortalisation of some of the available cultures^12^, which increases the possibility of performing more experiments with a given culture. We have used immortalised cultures to further increase this possibility.

Although we are concerned about the efficiency limitations of our gene editing protocol, specially due to transfection difficulties in myoblasts reported also by other laboratories^27,38^; we have successfully applied it to edit 2 different regions in two different cell backgrounds (Control and DMD), and we consider that those described and fully characterised in this manuscript could be relevant research models that we would be happy to share. The first of these models is an immortalised DMD disease cell culture model, (DMDΔ52-Model) that lacks exon 52 of the *DMD* gene, which disrupts the ORF and dystrophin expression. This model could be useful to evaluate mutation-independent drug treatments, and also exon skipping drugs that aim to skip exons 51 or 53^43,44^ as skipping either exon in this case would restore the ORF and dystrophin expression. We have demonstrated that DMDΔ52-Model lacks dystrophin expression and that this can be reverted through treatment with an exon 51 skipping drug. However, the main interest of this model was demonstrating the efficacy of the technique, as immortalised cultures from patients with the same deletion are currently available. We have replicated this protocol to generate other DMD-like cultures (details will be included in a manuscript being currently drafted).

An immortalised cell culture model constitutively expressing utrophin, DMD-UTRN-Model, is both a proof of principle of a possible therapeutic option to overexpress utrophin as a substitute for dystrophin, and a valuable research tool. The search of drugs that could overexpress utrophin is ongoing^45^ after the recent failure of ezutromid to show relevant results in clinical trials. Many research projects, including drug re-purposing screening, are ongoing to find new candidates to test in clinical trials^46^. However, there are no reliable positive controls that could be used to compare such treatments. We propose that our cell model could serve that purpose, offering researchers useful custom controls for their studies. We have tested this hypothesis and compared the stable utrophin overexpression quantified in the DMD-UTRN-Model with the utrophin that is expressed after treatment of the unedited DMD patient cultures with ezutromid, the lead market candidate in this field until very recently, with positive results. Previous studies in muscle sections show that DMD patients already overexpress utrophin, in many cases 4 to 5-fold the levels seen in control muscle sections^47^. Our choice to target this particular *UTR* region, increases basal overexpression in DMD cultures, and the amount of overexpression varies significantly when evaluated by western blot (more than 2.5 times) or our preferred method, myoblots (close to 1.5 times). We like to consider that myoblot evaluation reflects more closely the actual protein expression, as it is not subjected to many of the inherent problems of western blotting when evaluating very large proteins^48^. This is why we cannot comment yet on the differences in expression between our study and other published studies that also aimed to overexpress utrophin by gene editing, but which 1) targeted different promoter regions (*UTRN* A or *UTRN* B) of utrophin and 2) evaluated their results by western blot analysis^38^. We would be interested on studying this matter further to analyse the differences in utrophin expression when targeting different regions.

As a conclusion, we have created two new cell culture models that we have already used as efficient tools in our search for new therapies for DMD, after optimization of a gene editing method to be applied to myoblasts, a rather difficult target. We expect our experience to be useful to other muscle researchers and we are looking forward expanding those experiments.

## METHODS

### CRISPR/Cas9 tools

Specific sgRNA guides were designed using the online bioinformatics tool http://crispr.mit.edu^49^. Ten different guides (five before and five after the target region) were designed targeting exon 52 flanking regions in *DMD* gene and another ten targeting a repressor binding site in the UTR 3’ region of *UTRN* gene and selected according to their score number (Table 1). They were cloned into a plasmid containing Cas9 from *S. pyogenes* with 2A-*EGFP* pSpCas9 (BB)-2A-GFP (PX458) (Addgene plasmid # 48138, deposited by Feng Zhang). All sgRNAs were cloned using BbsI sites.

### Cell cultures

Human embryonic kidney 293 (HEK 293) cells, used in the preliminary selection of the best sgRNA combinations for our experiments, were purchased from the European Collection of Authenticated Cell cultures (ECCAC) via Sigma-Aldrich, Spain, and maintained following the manufacturer’s protocols.

Immortalized myoblasts derived from muscle biopsies from healthy controls and DMD patients were provided by the CNMD Biobank, London, UK and the Institut de Myologie Paris, France, and cultured using skeletal muscle medium (SMM) (Promocell, Germany) and differentiation media as previously described^48^.

### Cell culture transfection

All different sgRNAs combinations were transfected into HEK 293 cells using lipofectamine 2000^®^ (Thermo Fisher Scientific), according to the manufacturer’s protocol. Myoblasts seeded in 6 well plates at 70-80% confluence were transfected with 1.5ug of each plasmid with the most efficient guide RNA combination using ViaFect™ (Promega) transfection reagent (1:5 ratio).

### FACS workflow of GFP positive myoblasts

48 hours after transfection, myoblasts were trypsinised and collected for fluorescence activated cell sorting (FACS) at the Cell Analytics Facility (BD FACS Jazz) Achucarro Basque Center for Neuroscience - (Leioa, Spain). GFP-positive cells were seeded individually in 96 well plates for clonal selection. The first colonies were visible around 7 days post-sorting. Clones were expanded from single cell to near-confluence and expanded into larger well plates to be harvested 15-30 days post-sorting. Myoblasts often developed elongated and stressed shapes during this clonal expansion after single cell sorting. Harvested cultures were aliquoted: some aliquots were frozen for archival; others were pelleted for DNA analysis, while replicates were cultured further for characterization by immunocytochemistry, western blot and myoblots (see schematic workflow in Supplementary figure 2).

### Analysis of gene editing

DNA was extracted from cell pellets using a QIAamp^®^ DNA Mini Kit, Qiagen. PCR amplification targeting the edited regions was carried out using Taq DNA Polymerase (Recombinant), Invitrogen, under the following conditions: preheating 3’ 94°C, 25 cycles of 94° for 3’, 94° 20’’, 63° 20’’, 72° 1’ and 72° 5’ and DMD-Seq-D52-DOWN-F2 and DMD-Seq-D52-DOWN-R2 primers (see Suplementary Table 1). PCR products were resolved in 2% TAE-agarose gels and purified with QIAquick^®^ Gel extraction Kit, Qiagen. PCR amplicons corresponding to the expected length were analysed by Sanger sequencing at the sequencing platform of Biocruces Bizkaia Health Research Institute using DMD-Seq-D52-DOWN-F2 and DMD-Seq-D52-DOWN-R2 primers (see Supplementary Table 1).

### Off-target analysis of mutations in clonal lines

Potential off-target region loci of each sgRNA used were predicted using CRISPOR bioinformatics tool http://crispor.tefor.net/. The six most probable off-target sequences per guide (Table 2 and 3) were analysed in the edited clones using genomic PCR and Sanger sequencing. Primer sets flanking off-target sites and the corresponding internal primers used for Sanger sequencing are listed in Supplementary Table 2.

### Primary Antibodies

Anti-dystrophin: Dys1 (Leica Biosystems), Mandys1, Mandys106 (The MDA Monoclonal Antibody Resource) and anti-dystrophin antibody (Abcam 15277). Anti-utrophin: Mancho7 (The MDA Monoclonal Antibody Resource). Anti-myosin heavy chain (MF20: Developmental Studies Hybridoma Bank). Anti a-actinin (Sigma-Aldrich A7732).

### Immunostaining assays

Original and edited clones for objectives 1 (DMDΔ52-Model) and 2 (DMD-UTRN-Model) were cultured and immunostained for dystrophin or utrophin expression. Edited clones were seeded into chamber slides and treated with a MyoD virus, (Applied Biological Materials Inc, Canada) to facilitate differentiation into myotubes^50^. After seven days differentiating, samples were fixed with 4% PFA. Cultures were permeabilised with Triton 0.5% and then blocked for half an hour with BSA 2%. Afterwards, immunostaining was performed overnight at 4°C with the required antibodies. Primary antibodies used for dystrophin staining were a mix of Dys1, Mandys1 and Mandys106 at 1:100 dilution and for utrophin staining was Mancho 7 diluted at 1:50. The following day, after being washed with PBS Tween 0.1%, cells were stained with Alexa Fluor 488 goat anti-mouse (Invitrogen) for 1 hour at room temperature. Hoechst 1/2000 was used for nuclei staining and chamber slides were mounted with PermaFluor^™^ Aqueous Mounting Medium (Thermoscientific). Images were captured with a LEICA DMI 6000B microscope at the Microscopy Platform at Biocruces Bizkaia Health Research Institute.

### In-cell western assay (myoblots)

Myoblots were performed as described before^48^. In short, clones were seeded in 96-well plates and incubated for 24 h in SMMC, after which they were treated with MyoD virus and incubated in differentiation media for 7 days. Then, plates were fixed with ice-cold methanol, permeabilised and blocked before incubation with the required primary antibodies overnight: anti-dystrophin mix (Dys1, Mandys1 and Mandys106 at 1:100), anti-utrophin (Mancho 7 antibody at 1:50), and anti-myosin heavy chain antibody (MF20 at 1:100) that was used to evaluate differentiation. Next day, plates were incubated with the secondary antibodies. Biotin-mediated amplification (Abcam 6788 goat antimouse IgG biotin 1:2000) was used to increase dystrophin signal. Secondary antibodies, IRDye 800cw streptavidin 1:2000 and IRDye 800CW goat anti-mouse 1:500, were prepared together with CellTag 700 Stain (LI-COR^®^ Biosciences) at 1:1000 dilution and incubated for 1 hour at RT and protected from light. After incubation, plates were analysed using the Odyssey^®^ CLx Imager (LI-COR^®^ Biosciences).

### Treatment with antisense exon skipping drugs

Cultures in 96 wells and P6 wells were treated with a 2’MOE-phosphorotioate antisense oligonucleotide (AO) aiming to skip DMD exon 51 (51-[T*C*A*A*G*G*A*A*G*A*T*G*G*C*A*T*T*T*C*T]-3⍰, Eurogentec, Belgium) by transfection with Lipofectamine as described in^43,44^ and analysed by either myoblot (96 well plates) or RT-PCR (pellets extracted from 6 well plates).

### RT-PCR

RNA was extracted from cell pellets (RNeasy mini kit, Qiagen) according to the manufacturer’s protocol. Reverse transcription of the samples was performed using (SuperScrip™ IV Reverse Transcriptase, Invitrogen) according to the manufacturer’s protocol. cDNA samples were either used for digital droplet PCR analysis or amplified by nested PCR using specific primers sets (Supplementary Table 1) and Taq DNA Polymerase (Recombinant), Invitrogen, as described in^44^ for exon skipping analysis. Amplified samples were resolved in TAE-agarose and PCR amplicons of interest were first analysed with Gel Doc TM EZ Imager, BIORAD and then purified with (QIAquick^®^ Gel extraction Kit, QIAGEN) for sequencing analysis. Before DNA extractions bands were semi-quantified using Image J.

### Treatment with utrophin overexpression drugs

Ezutromid was diluted first in DMSO and finally in differentiation medium to different concentrations and added to myoblasts in 96 well plates 7 days after differentiation. Twenty-four hours after treatment, medium was removed, and plates were fixed with ice-cold methanol for myoblot analysis.

### Western blot

Cell cultures were seeded into P6 plates and trypsinized after 7 days of differentiation. Then, cell pellets were solubilized in lysis/loading buffer and denatured at 95°C for 5 min. Samples were loaded onto a NuPAGE^®^ Novex^®^ 3–8% Tris-Acetate Gel3–8% (Thermo Fisher Scientific) and run in Novex Tris-Acetate SDS Running Buffer (Thermo Fisher Scientific) for 60 min at 70 V + 120 min at 150 V at 4°C. Protein wet transfer was performed overnight at 4°C using an Immobilon^®^-FL PVDF membrane (Merck™). Next day, membranes were stained with Revert ™ 700 Total Protein Stain (Li-Cor) for total protein measurement, blocked with Intercept (PBS) Blocking Buffer (Li-Cor) for 2 hours and incubated overnight at 4°C with the primary antibodies (1/200 anti-dystrophin antibody Abcam15277, 1/50 anti-utrophin antibody Mancho 7 or 1/500 anti a-actinin antibody, Sigma-Aldrich A7732). After washing steps with PBS-Tween 0.1%, membranes were incubated with secondary antibodies for 1 hour (1/5000 IRDye 800CW goat anti-rabbit 926–32211 or IRDye 680RD goat anti-mouse 926-68070, Li-Cor) at room temperature, washed again and scanned using an Odyssey Clx imaging system. Bands quantification was performed using Image Studio ™ software.

### Droplet digital PCR (ddPCR)

Gene expression was detected and quantified using a QX200™ Droplet Digital™ PCR system (Bio-Rad). The reaction was performed using 2 μl of cDNA in a 20 μl reaction volume containing: 1 μl of Utrophin Taqman probe (ID: Hs01125975, FAM labelled), 1μl of MYF5 ddPCR™ GEX Assay probe (ID: dHsaCPE5026295, HEX labelled) or MYH2 ddPCR™ GEX Assay probe (ID: dHsaCPE5050991, HEX labelled), 10 μl of ddPCR™ Supermix for Probes (no dUTP) (Bio-Rad) and 6 μl of DNase/RNase-free H2O.

To generate the droplets, 20 μL of the previous ddPCR reaction and 70 μL of Droplet Generation Oil for Probes (Bio-Rad) were added to the 8 channel droplet generation cartridge according to manufacturer instructions and this cartridge was placed in the QX200 droplet generator (Bio-Rad). Then, 40 μL of the resulting droplet emulsion was transferred to a semi-skirted 96 well PCR plate (Eppendorf), sealed with foil and amplified on a thermal cycler using the following amplification conditions: enzyme activation 10’, 40 cycles of 94°C for 30’’ and 55°C for 1’, and heat deactivation 10’ 98°C. Plates containing the amplified droplets were loaded into the QX200 droplet reader and results were analysed using QuantaSoft software™ (Bio-Rad).

### Statistical analysis

Mann-Whitney U test was used throughout this study to calculate P values for determination of statistical significance (*p-value<0.05, ** p-value<0.01, ****p-value<0.0001). Statistical analysis was performed using GraphPad Prism software.

## ACKNOWLEDGEMENTS

We acknowledge the use of cell cultures provided by the Queen Square Centre for Neuromuscular Disorders BioBank (CNMD Biobank, London, UK) and the Institut de Myologie (Paris, France, immortalised cultures).

We gratefully acknowledge the use of the antibodies provided by Professor Glenn Morris from the Muscular Dystrophy Association (MDA) Monoclonal Antibody Resource, which distributes free antibodies for research in neuromuscular diseases worldwide from Oswestry, United Kingdom.

The MF20 antibody developed by D.A. Fischman, Weill Cornell Medical College, was obtained from the Developmental Studies Hybridoma Bank, created by the NICHD of the NIH and maintained at The University of Iowa, Department of Biology, Iowa City, IA 52242.

Cell sorting experiments were performed at the Cell Analytics Facility at Achucarro-Basque Centre of Neuroscience (Leioa, Spain).

We would like to thank Ms. Karmele Alapont-Celaya for her assistance in the completion of some laboratory tasks during this project. Drs. Gustavo Pérez-Nanclares and Ana Belén de la Hoz at the Genetics-Genomics Facility, Biocruces Bizkaia Health Research Institute, are also gratefully acknowledged for their help with sequencing reactions. Special thanks are due to Drs. Maria Dolores-García Vázquez at the Cell Culture Facility and Dr Javier Díez-García at the Microscopy Facility, both at Biocruces Bizkaia Health Research Institute, for their assistance at their corresponding platforms.

## DATA AVAILABILITY

Under request

## AUTHOR CONTRIBUTIONS STATEMENT

P S-M, formal analysis, investigation, data curation, writing or original draft, review and editing, visualization, supervision. E. A-A, investigation, data curation, writing - review and editing. A. A-M, investigation, writing - review and editing. L. d-l-P-O, investigation, writing - review and editing. I. G-J, methodology, investigation, review and editing, supervision. G. G-I, investigation, writing - review and editing. I. L-A, investigation, writing - review and editing. A. L-M, investigation, writing - review and editing. J. P-G, investigation, writing - review and editing. E. R-D-Y, methodology, investigation, writing - review and editing, supervision. F. G conceptualization, formal analysis, writing - review and editing. V. A-G: conceptualization, methodology, formal analysis, investigation, resources, data curation, writing or original draft, review and editing, visualization, supervision, project administration and funding acquisition.

