## Supplementary data for "Duchenne Muscular Dystrophy Cell Culture Models Created By CRISPR/Cas 9 Gene Editing And Their Application To Drug Screening"

| NAME | PRIMERS SEQUENCE | APLICATION | TARGET | AMPLICON LENGTH (bp) (NON EDITED OR NON SKIPPED) | AMPLICON LENGTH (bp) (EDITED OR SKIPPED) |
|---|---|---|---|---|---|
| **DMD-Seq-D52-DOWN-F2** | TTTCTAAAAGTGTTTTGGCTGGTC | Editing confirmation | DNA | 750 | 228 |
| **DMD-Seq-D52-DOWN-R2** | TACCAAAGTTCCTGCCCACC | Editing confirmation | DNA | 750 | 228 |
| **UTRN F1** | TGATGGTACCTCCACCTACATCT | Editing confirmation | DNA | 692 | 421 |
| **UTRN R1** | TTACTTCCCATTGTTACTGCAA | Editing confirmation | DNA | 692 | 421 |
| **47F** | AGTGCTCCCATAAGCCCAGAA | Skipping confirmation | RNA | 1265 | 1147 |
| **54R** | GAAGTTTCAGGGCCAAGTCA | Skipping confirmation | RNA | 1147 | 914 |
| **49F** | CCAGCCACTCAGCCAGTG | Skipping confirmation | RNA | 1147 | 914 |
| **53R** | TTGCCTCCGGTTCTGAAGG | Skipping confirmation | RNA | 422 | 189 |

**Supplementary table 1. Gene edition and antisense experiments primer sets.**

| Primers off targets (5'—3'): | |
|---|---|
| **Ob1_2_Off1_F** | ATGCTTCTCATTTGCTGCCTGATG |
| **Ob1_2_Off1_R** | GCTGTCACGATCTGATTGGAGTTTC |
| **Ob1_2_Off1_Seq** | TCTCCAGGCTCCAAGTAT |
| **Ob1_2_Off2_F** | ACAAGAAAGTCCTGGGTGTCA |
| **Ob1_2_Off2_R** | GACTTGTCAGCCTCCATAATCATT |
| **Ob1_2_Off2_Seq** | CTGATGAGGGCTTTCCAG |
| **Ob1_2_Off3_F** | AATGGCAGTTTTGGGAAATTCA |
| **Ob1_2_Off3_R** | CATTTTTTGTTGTCTGTGCCC |
| **Ob1_2_Off3_Seq** | TAAGTTACAATGCTTCCTGAA |
| **Ob1_2_Off4_F** | TACTTAATCCCCGTGTGTCTCAGT |
| **Ob1_2_Off4_R** | ATGCAATGTGAAGGCTGTCCG |
| **Ob1_2_Off4_Seq** | TTCCTTGTTTGTGCCTCA |
| **Ob1_2_Off5_F** | TCAAGAAAGTATGGTGTTGGTGAA |
| **Ob1_2_Off5_R** | AATTGTGTGAGTCCATTCCCATAA |
| **Ob1_2_Off5_Seq** | TTGACAAAGGGGCAAAGG |
| **Ob1_2_Off6_F** | ATGGCAAATAGTTACAATGTCA |
| **Ob1_2_Off6_R** | AATTTACACAACTAGAGCCATCT |
| **Ob1_2_Off6_Seq** | ACAGGAAGTCTTGAAAAAAGTA |
| **Ob1_6_Off1_F** | TTCCCTTGAACTTGCGGAGGT |
| **Ob1_6_Off1_R** | CCTTTTGGTTCCTGTGCCCC |
| **Ob1_6_Off1_Seq** | GTGCTCAAGCTCCCTTAT |
| **Ob1_6_Off2_F** | GCAGCAGATGGGTTAGGAGG |
| **Ob1_6_Off2_Seq** | CATGTAGGAATGACAGGAGT |
| **Ob1_6_Off2_R** | CTTGTGGCCAGCATCAGGTA |
| **Ob1_6_Off3_F** | CTGCTAGAGTGAGAGAACTGTGG |
| **Ob1_6_Off3_R** | GGAAACCAGGGCAAATCATGTCT |
| **Ob1_6_Off3_Seq** | GCCCCACATAGGACAAAT |
| **Ob1_6_Off4_F** | GCTGAAGGAAGTTCCAGGCA |
| **Ob1_6_Off4_R** | CAGGCTGGCAAGATGGAGAA |
| **Ob1_6_Off4_Seq** | GGCATCCTTATAGCAATTTTT |
| **Ob1_6_Off5_F** | GATAGCCCCACCAGACAACC |
| **Ob1_6_Off5_R** | AGGGCCAAATCCTCACAACC |
| **Ob1_6_Off5_Seq** | TCACCATTCTCATCCCCT |
| **Ob1_6_Off6_F** | TCAGAAAGGCTTGCCCTCA |
| **Ob1_6_Off6_R** | CTCACATGGCAAGTGGGGAT |
| **Ob1_6_Off6_Seq** | CCATCTAAAATCACCACACC |
| **Ob2_22_Off1_F** | ATGAGCCTCACAGATGCCTG |
| **Ob2_22_Off1_R** | GAAGACAGGGCCTGGATGTC |
| **Ob2_22_Off1_Seq** | TTAAAGTCTGTGCCCCTC |
| **Ob2_22_Off2_F** | AGGCTCTGCAGTTCAACCTC |
| **Ob2_22_Off2_R** | AACAGGCTCCAAACGTGTGA |
| **Ob2_22_Off2_Seq** | AATGACTTATACAGGGGACAT |
| **Ob2_22_Off3_F** | GGCCTTCTTTCGTGGACAGA |
| **Ob2_22_Off3_R** | GAATCATAGGCCTCCCGTGG |
| **Ob2_22_Off3_Seq** | TCGTGTTCTCTGTTGTGA |

| | |
|---|---|
| **Ob2_22_Off4_F** | GCCATAATCACACATCAAACCCT |
| **Ob2_22_Off4_R** | TGTTGCCATGCGAATTCGAG |
| **Ob2_22_Off4_Seq** | GAGAAGTTGAGGGAACCG |
| **Ob2_22_Off5_F** | GAGGTGGCATTCGGTAAAAGTTC |
| **Ob2_22_Off5_R** | TTGACAACCACGGGAGGCAG |
| **Ob2_22_Off5_Seq** | TGATTCTTTCCAGGCTCAT |
| **Ob2_22_Off6_F** | GGAGTGTGAGGGCTTCCTTC |
| **Ob2_22_Off6_R** | AGATGCCTGCTTACCTGCTG |
| **Ob2_22_Off6_Seq** | CATCTGTGTTCTCTGTTGTG |
| **Ob2_26_Off1_F** | GGGCAGGCTTGGGAGACATA |
| **Ob2_26_Off1_R** | GTGTCCAGCCCATTCTTTGAAGT |
| **Ob2_26_Off1_Seq** | TGCTCCCACTGCTGTTAG |
| **Ob2_26_Off2_F** | AGGTGCTCGCTTCTTTCCAA |
| **Ob2_26_Off2_R** | CCAGAAGTGAAGCTTTGCACC |
| **Ob2_26_Off2_Seq** | AACTTCTTGCACAGCCTT |
| **Ob2_26_Off3_F** | GGCATTCCTAGATCAGTGTGTGC |
| **Ob2_26_Off3_R** | CCCCAACTCAAACCAAGACGG |
| **Ob2_26_Off3_Seq** | CTGGGGGGATGTTACTGT |
| **Ob2_26_Off4_F** | TGGCTCTGTTTCTTGCCCAA |
| **Ob2_26_Off4_R** | TGGTTACTGGGCAGACATGG |
| **Ob2_26_Off4_Seq** | AAGCTAAAAGACATTGACAGT |
| **Ob2_26_Off5_F** | CACTGGAAAGAACATGGGCTCTG |
| **Ob2_26_Off5_R** | TGGTGTGTCCTGGGAGCATC |
| **Ob2_26_Off5_Seq** | CTCGGTTTCTACTGTGTGA |
| **Ob2_26_Off6_F** | TTTTATGTCCCCCACCCCTCA |
| **Ob2_26_Off6_R** | CCATGCCCAGCCCTAGTTTG |
| **Ob2_26_Off6_Seq** | GGGAAGCAGAGAAGTTGT |

**Supplementary table 2. Off-target primer sets.**

**Figures**

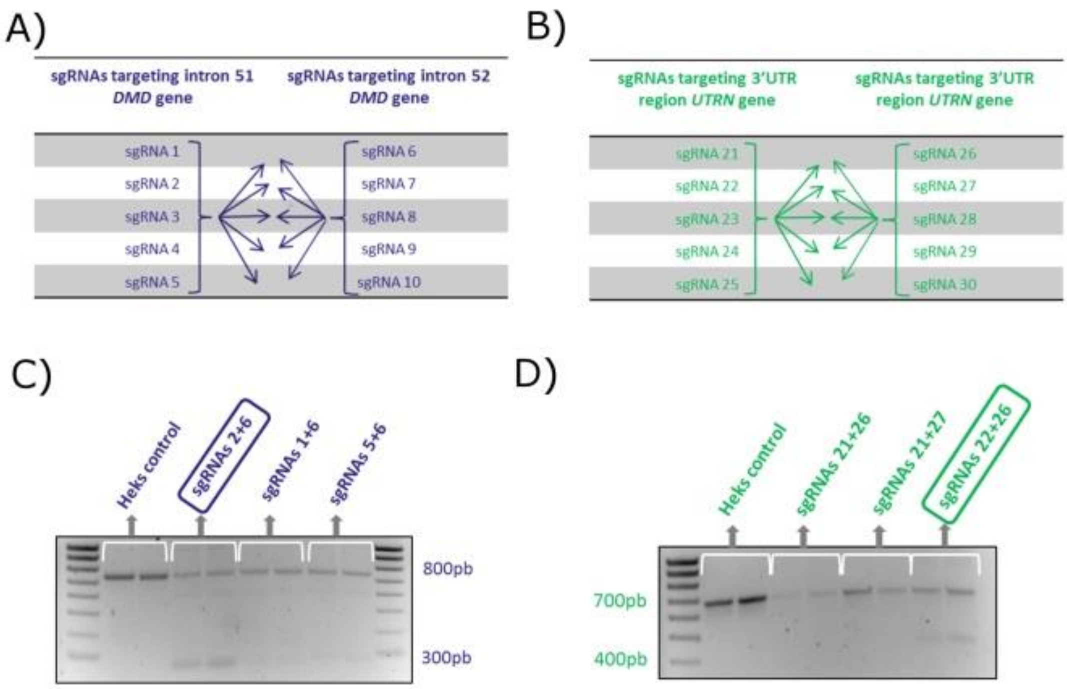


**Supplementary figure 1. sgRNAs pairs test in HEK293 cells.**

(A and B). Representation of all the different sgRNAs combinations tested for editing the *DMD* (A) and the *UTRN* loci (B). (C and D) Representative PCR analysis of HEK293 cells transfected with some of the sgRNAs combinations tested. Upper bands correspond to wild type or non-edited cells, while the lower bands correspond to the edited ones. Samples were analysed in duplicates (marked in white). Selected sgRNAs combinations are highlighted: Obj1sgRNA2 (C); Obj2 sgRNA22+sgRNA26 (D).

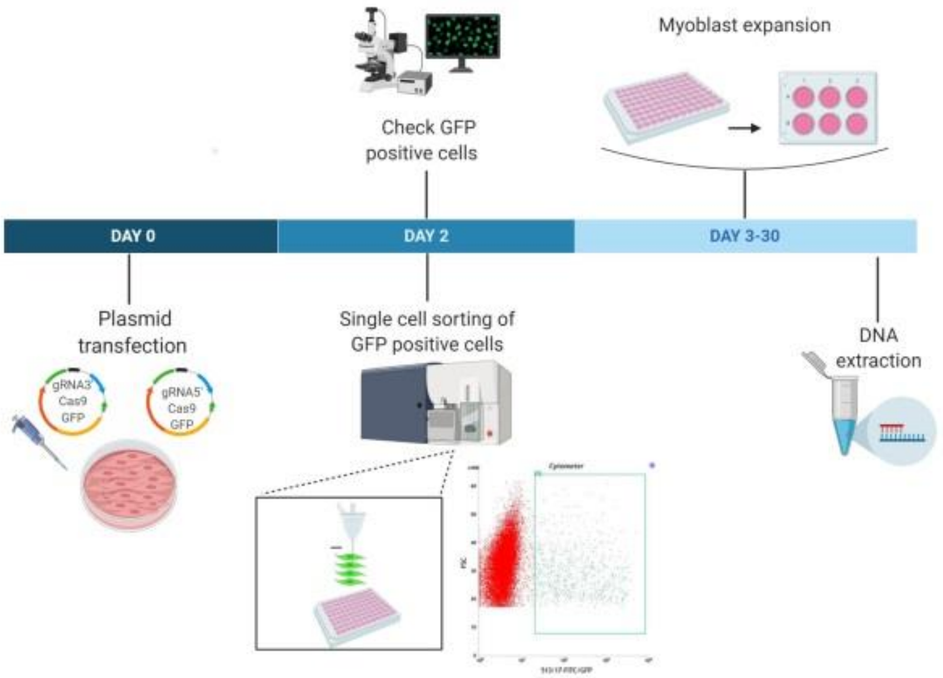


**Supplementary figure 2. Cloning and editing workflow diagram.**

Scheme of the workflow followed to obtain the edited myoblast cell lines. 48 hours after plasmid transfection GFP positive myoblasts were single cell sorted using FACS. Clones were expanded until confluence for DNA extraction. The dot plot shows GFP positive myoblasts (3,22 %) selected using FACS 48h post transfection with ViaFect™ reagent. Created with BioRender.com

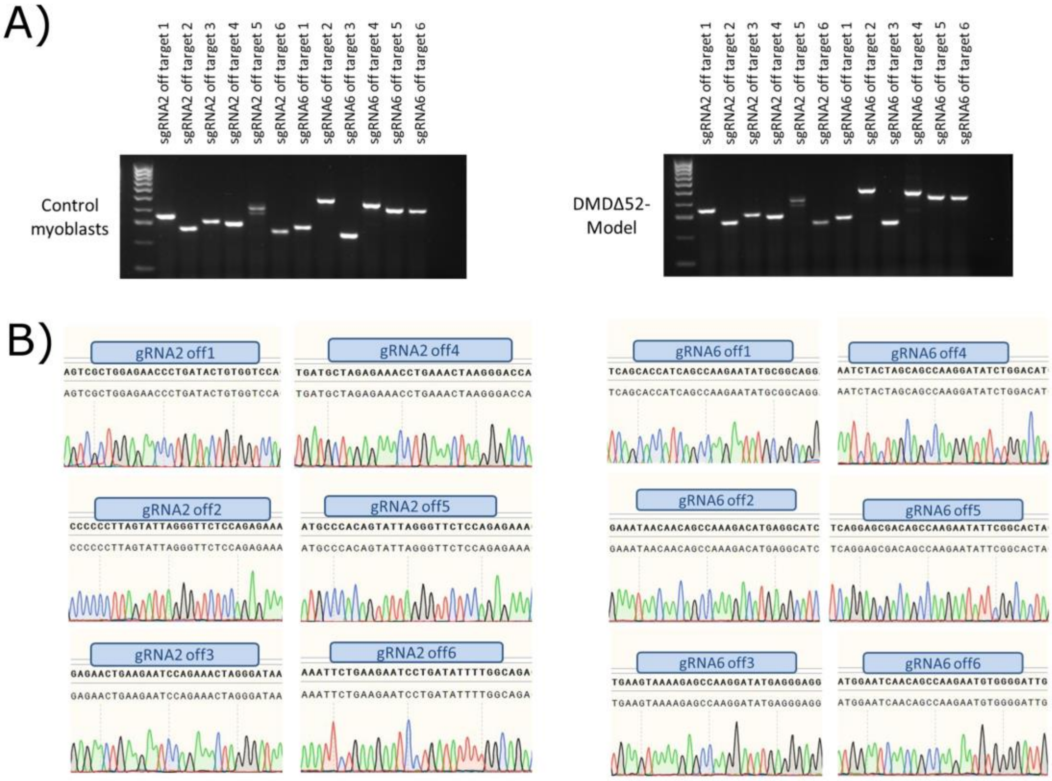


**Supplementary figure 3. PCR Analysis of Off-target Effects. Representative gel and sequence of PCR analysis performed for all targets.**

(A) Agarose gel showing the amplification of the six predicted off-targets regions for sgRNA2 and the six for sgRNA6 (the combination used for our DMD edition model) amplified in control myoblasts and DMDΔ52-Model. (B) All the amplicons were sequenced and no differences between control myoblasts and DMDΔ52-Model were found.

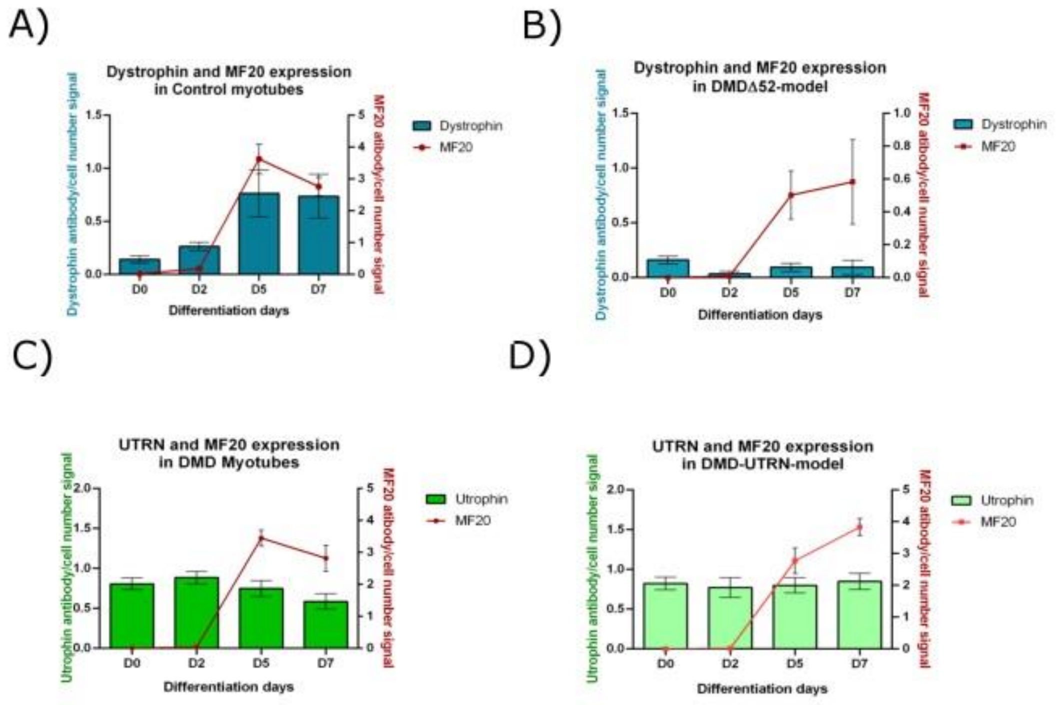


**Supplementary figure 4. Myoblot assays at different fusion times in edited cells.**

(A and B) Dystrophin and MF20 expression determined by myoblot in DMDΔ52-Model compared to control myoblasts along differentiation process. (C and D) Utrophin and MF20 expression determined by myoblot in DMD-UTRN-Model compared to DMD myoblasts. Myoblot analysis was performed using n=24 replicate wells for dystrophin or utrophin staining and n=18 replicate wells for MF20 staining.
